# The transcriptional response of genes to RpoS concentration in *Escherichia coli* is not determined by core promoter sequences

**DOI:** 10.1101/796656

**Authors:** Suzannah M. Beeler, Christopher R. Doering, Sarena Tran, Daniel M. Stoebel

**Author notes:** **Corresponding author:** Daniel Stoebel.

## Abstract

The alternative sigma factor RpoS is an important regulatory protein in *Escherichia coli*, responsible for mediating the general stress response. RpoS levels vary continuously in response to different stresses. Previous work has shown that genes vary in their responsiveness to increasing RpoS concentrations, with some genes being “sensitive,” requiring only a low level of RpoS to be relatively highly expressed, while other genes are “insensitive,” only being highly expressed in the presence of high levels of RpoS. In other systems, this type of variation is caused by interactions between the regulatory protein and the DNA it binds. To see if this is the case for RpoS, we measured twelve RpoS binding site mutants for their effects on maximal expression and responsiveness to increasing RpoS concentration. While maximal expression varied over an order of magnitude across these twelve constructs, the responsiveness to increasing RpoS concentration was largely unaffected, suggesting that the RpoS binding site alone is not responsible for a genes’ sensitivity or insensitivity to RpoS. In addition, we swapped the RpoS binding region between sensitive and insensitive promoters and found no change in the behavior of the promoter. Taken together, these results argue that differences in sensitivity of the RpoS-dependent promoters are not due to interactions between RpoS and the various DNA sites it binds.

## INTRODUCTION

Transcription in bacteria requires sigma factors that bind to RNA polymerase (RNAP) and help enable promoter binding and transcription initiation (Borukhov and Severinov, 2002). *Escherichia coli* has seven sigma factors, each of which regulates a particular suite of genes (Gruber and Gross, 2003). For example, RpoD (also known as σ^70^ or σ^D^) is known as the housekeeping sigma factor as it is essential for survival and is responsible for transcribing genes needed for cell growth. RpoS (also known as σ^38^ or σ^S^) is responsible for the general stress response and regulates genes involved in responding to stressors like cold shock, acid stress, osmotic stress, and entry into stationary phase (Battesti et al., 2011).

Since the genes in the RpoS regulon are only needed in the presence of a stressor, RpoS is tightly regulated to keep the expression of stress response genes low unless necessary (Battesti et al., 2011). This regulation of RpoS occurs at the level of transcription, translation, protein degradation, and protein activity (Battesti et al., 2011; Gottesman, 2019; Hengge, 2009; Lange and Hengge-Aronis, 1994). This regulation results in only low levels of RpoS while *E. coli* K-12 is in exponential growth in rich media at 37 °C. However, as a culture reaches stationary phase or is faced with some other stressor (like cold-shock or increased osmolarity), the level of RpoS begins to increase, allowing the cells to better cope with this stress (Battesti et al., 2011; Lange and Hengge-Aronis, 1994; Schellhorn, 2014). Changing regulation of RpoS expression during the transition from exponential growth to stationary phase results in a continuous rise of RpoS levels during this stress response (Lange and Hengge-Aronis, 1994).

The continuous nature of possible RpoS levels has important consequences for the RpoS regulon. We recently used RNA-seq to show that members of the RpoS regulon respond differently to changes in RpoS level (Wong et al., 2017). In particular, we found that some genes are sensitive to increasing RpoS levels (reaching near maximal expression at low RpoS levels, such as *astC*), while other genes are insensitive (requiring a high level of RpoS to be maximally expressed, such as *gadB*). Genes with these different expression patterns have different physiological functions and appear to differ in the timing of their expression in response to the onset of stationary phase (Wong et al., 2017). Differences in the response to RpoS level likely coordinate patterns of transcription in response to stresses.

The mechanistic basis of this difference in response to RpoS levels is unclear. In the cases of Spo0A and CodY in *Bacillus subtilis* and PhoB and LexA in *E. coli*, interactions between the regulatory protein and its DNA binding site in the promoter determines the level of the protein required for induction (Brinsmade et al., 2014; Culyba et al., 2018; Fujita et al., 2005; Gao and Stock, 2015). In addition, consideration of the basic biochemistry of transcription can provide intuition of how RpoS level might influence transcriptional output. If the RNAP-σ^38^ complex binds to these core promoters with simple Michaelis-Menten kinetics (Brewster et al., 2012; Újvári and Martin, 1996), then we could expect to see response curves that vary from a nearly switch-like behavior (when the binding affinity is high) to something more gradually increasing (when binding affinity is low), explaining much of the variation in promoter response to RpoS level we previously observed. By examining the response of different core promoters individually as well as in the context of different whole native promoters, we can tease apart the relative effects of the core promoter and additional regulation in determining the response to increasing RpoS.

## MATERIALS AND METHODS

### Strains and Growth Conditions

The strains used for this study are listed in Table S1. Unless otherwise noted, cultures were grown aerobically (at 225 rpm) in 5 mL of LB (0.5% yeast extract, 1% tryptone, 1% NaCl) at 37°C in vertical 16 × 150 mm test tubes. Where necessary, cultures were grown with ampicillin at 100 µg/mL for plasmids or 25 µg/mL for chromosomal copies.

### Strain creation

Promoters for plasmids pST1 – pST17 (Table S2) were created by synthesis of oligonucleotides that yielded the desired double stranded substrate when annealed (Table S3). These constructs were flanked with KpnI and EcoRI cut sites to allow for ligation into pLFX. To make the double stranded RpoS binding site region, 1 µM of forward and reverse oligos were heated for one minute at 100°C with 5 mM MgCl_2_ and 7 mM Tris-Cl (i.e. Qiagen Elution Buffer) and annealed by slowly cooling to room temperature.

Cloning of promoters pST1 – pST17 into pLFX was achieved by digesting both the annealed promoter constructs and pLFX with EcoRI-HF and KpnI-HF for 30 mins at 37°C, followed by dephosphorylation with Antarctic Phosphatase for 1 hour at 37°C. The digests were then purified with either GenElute PCR Clean-Up Kit (Sigma-Aldrich) or QIAquick PCR Purification Kit (Qiagen), followed by ligation with T7 ligase (New England Biolabs) for 30 mins at 25°C. 5 µL of ligated plasmid was then transformed into competent BW23473 cells (made using the *Mix & Go E. coli* Transformation Kit & Buffer Set, Zymo Research) and plated on LB + amp plates and grown overnight. Possible transformant colonies were inoculated in LB + amp and grown overnight. Plasmids were isolated in a 3 mL prep using Zyppy Plasmid Miniprep kit (Zymo Research), and inserts were verified by using Sanger sequencing.

Promoters with mutations in the -10 region (plasmids pDMS163 – pDMS168; Table S2), and the core promoter swaps (pDMS213 and pDMS217; Table S2) were created by site-directed mutagenesis using the Q5 site-directed mutagenesis kit (New England BioLabs). For -10 region mutations, primer pairs (table S3) were used to amplify pST1 using the manufacturers suggested reagent concentrations. For core promoter swaps, primer pairs amplified pDMS157 and pDMS160 as template. PCR was performed with an initial denaturation of 98° C for 30 s, followed by 25 cycles of 98° C for 10 s, 58° C for 30s, and 72° C for 3 min. PCR concluded with a final extension of 72° C for 3 min. PCR was followed by the kinase, ligase, and DpnI treatment steps according to the manufacturer’s recommendations. Cells were transformed into chemically competent BW23473 cells and plated on LB + ampicillin. Transformants were miniprepped and inserts were verified by Sanger sequencing.

Fusion plasmids were integrated into strain DMS2564 (Wong et al., 2017) with helper plasmid pPFINT (Edwards et al., 2011) and single-copy integrants were confirmed using the PCR assay of Haldimann and Wanner (Haldimann and Wanner, 2001).

### β-galactosidase assays

Strains were grown for 20 hours at 37°C with 0%, 10^−4^%, and 10^−2^% arabinose to yield RpoS concentrations of 0%, ∼26%, and ∼89% of wild type, respectively (Wong et al., 2017). β-galactosidase levels were measured using the method of Miller (1992). A 96-well plate spectrophotometer (BioTek) was used for measurements, so Miller unit values reported here cannot be directly compared to those taken with individual 1cm cuvettes.

### Data analysis

Sensitivity of a promoter was quantified as in Wong et al. (2017). Briefly, for each replicate we calculated the distance between the observed expression at the intermediate RpoS concentration and the expected level based on a linear pattern, standardized by the difference in expression between high and low RpoS conditions. Statistical testing of changes in sensitivity was performed with a two-sample randomization test.

All data analysis was performed in R (R Core Team, 2018).

## RESULTS

To investigate the role that RpoS-dependent core promoters play in determining the responsiveness of gene expression to varying RpoS concentrations, we began by creating and testing a total of twelve constructs (Table 1). From the consensus promoter (Typas et al., 2007; Wong et al., 2017), we introduced several types of mutations: increased GC content, altered spacer length between the -10 and -35 binding sites, and mutated residues in the -10 binding site. These mutations were expected to influence transcriptional initiation to different extents and by different mechanisms, whether by altering the ability of RNAP-σ^38^ to bind to the promoter or by making DNA melting and subsequent initiation more difficult. In our previous ChIP-seq work we were unable to find a consensus motif for the -35 region (Wong et al., 2017); mutations targeting that region were not constructed.

**Table 1.**
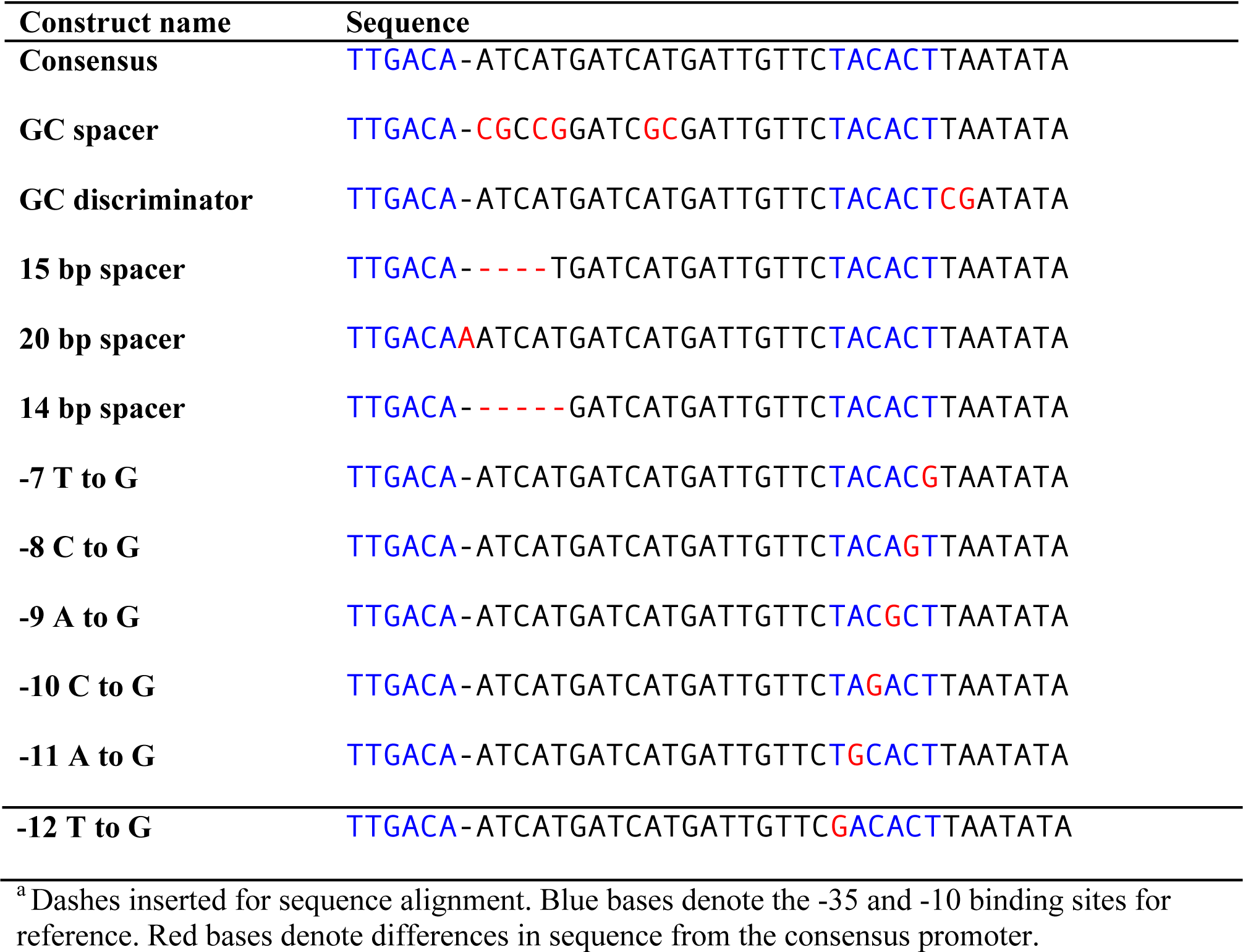
RpoS binding sites tested^a^.

We measured the extent to which the 12 promoters drove expression of *lacZ* in the presence of three RpoS concentrations (0%, 26%, and 89% of wildtype expression) using β-galactosidase assays. There is about a 10-fold change in maximal expression across the twelve constructs, from 328 ± 5 Miller units (consensus sequence, mean ± SE) to 35 ± 1 Miller units (−11 A to G single basepair substitution) (Figure 1). As expected, the consensus sequence had the highest activity of all 12 constructs.

**Figure 1.**
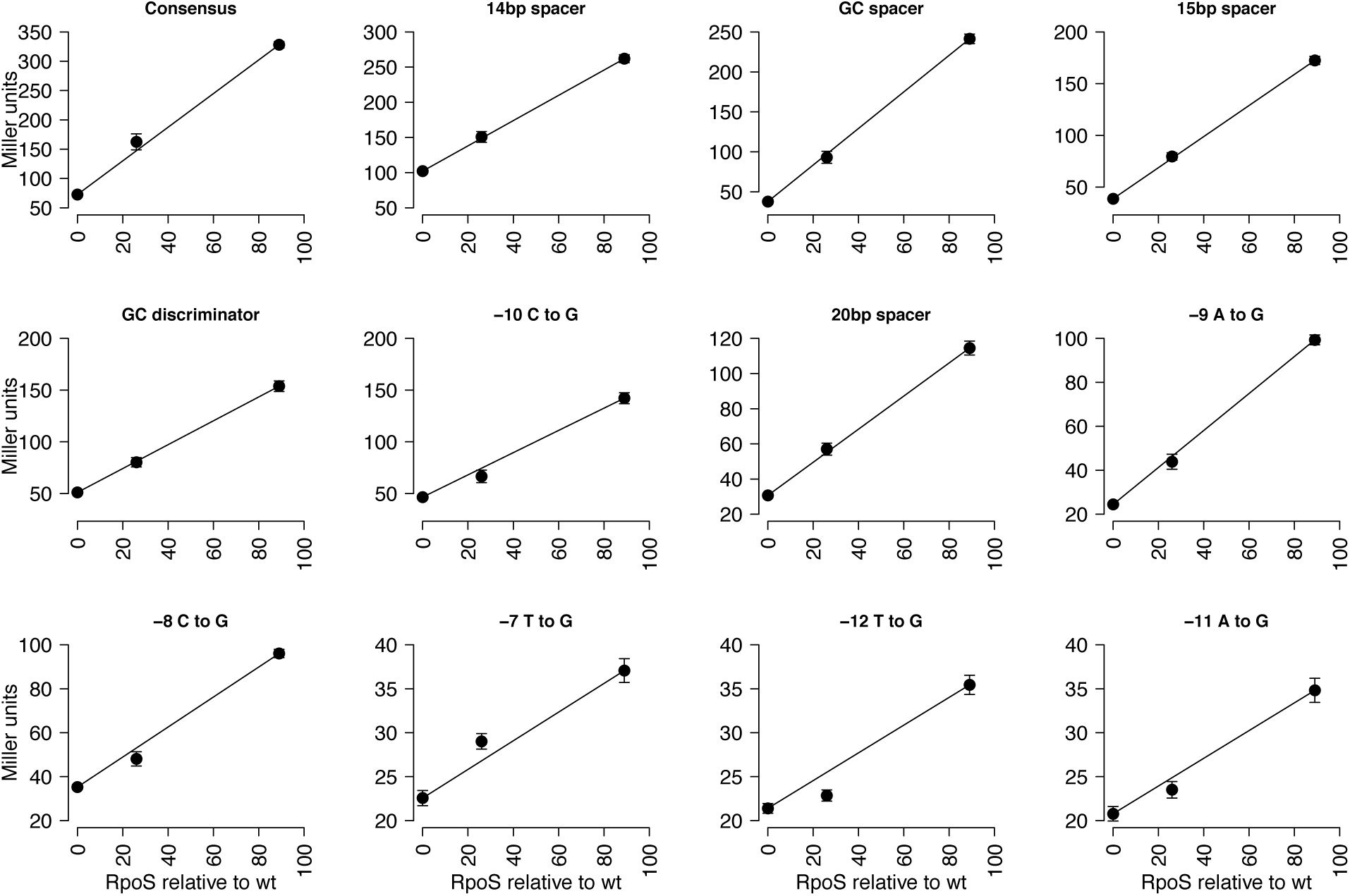
Expression patterns of the various RpoS binding site constructs over varying RpoS concentrations as measured by β-galactosidase assay. Constructs are ordered by maximal expression, with highest in the upper-left and lowest in the lower-right. n = 7 – 8, error bars represent standard error of the mean.

While the 12 constructs vary in maximal expression 10-fold, they all show a largely linear response to RpoS levels. None of mutant promoters differ significantly in their sensitivity from the consensus promoter (p > 0.05, two-sample randomization test, 100,000 replicates; p-values adjusted by the method of Holm (1979)). Based on work in other systems (Brinsmade et al., 2014; Culyba et al., 2018; Fujita et al., 2005; Gao and Stock, 2015), we expected there to be a positive correlation between the maximal activity of a promoter and the sensitivity. However, there was no significant correlation between the maximal expression and the sensitivity of each construct (r = 0.33, p = 0.3; Figure 2). Sensitivity values varied within a narrow range (−0.17 to 0.16), a small part of the possible variation, and the variation seen in naturally occurring promoters. For example, the wild type *gadB* promoter has a sensitivity of −0.25, and the wild type *astC* promoter has a sensitivity of 0.68 (Figure 3).

**Figure 2.**
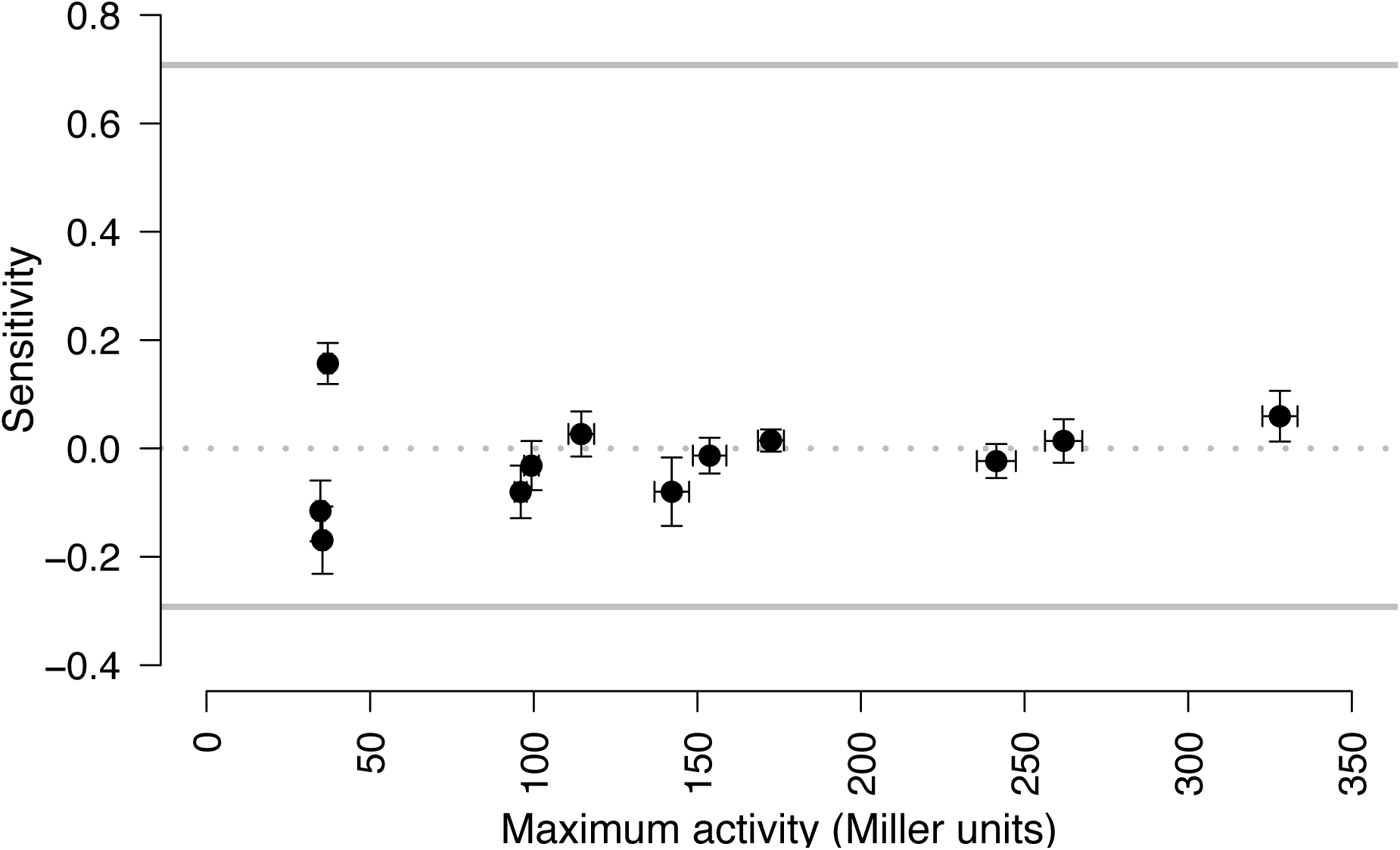
Sensitivity and maximal activity of each of the twelve constructs. n = 7 – 8, and error bars represent standard error of the mean. The correlation between the two variables is not significant (r = 0.33, p = 0.3). The upper and lower gray lines represent the maximum and minimum possible values for sensitivity with a monotonic response to RpoS. The dashed gray line represents sensitivity of 0.

**Figure 3.**
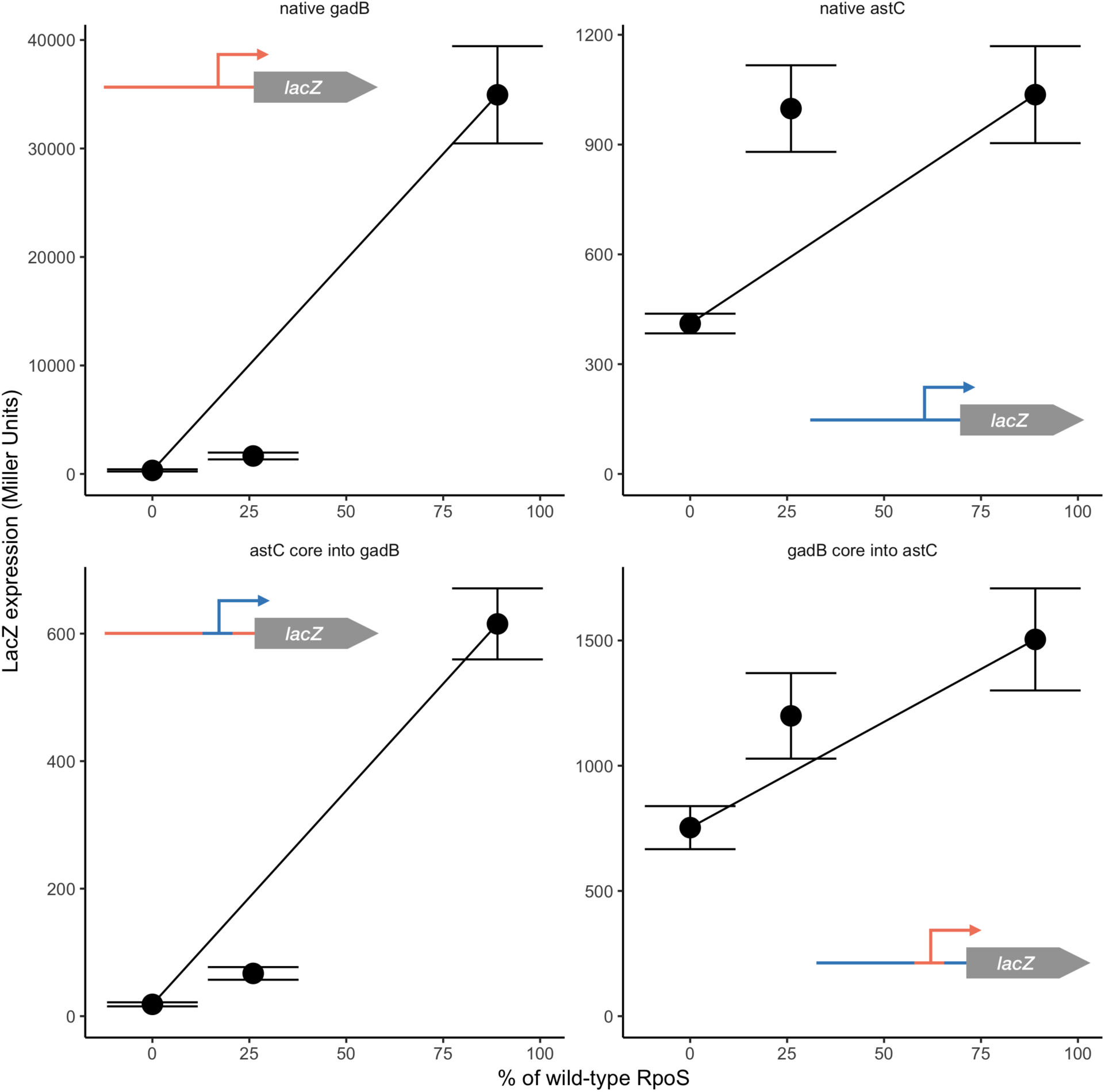
Effect of swapping core promoters from *gadB* and *astC* into full-length promoters. *astC* moved into the full length *gadB* promoter is insensitive, just as the native *gadB* promoter is. The *gadB* core promoter moved into the *astC* is sensitive, as is the native *astC*. The *astC* into *gadB* promoter is slightly less sensitive than the native *gadB*, a significant difference (p = 0.02, two-sample randomization test), while the swapped full-length *astC* is not different from native *astC* (p = 0.41, two-sample randomization test).

### Core promoters in their natural contexts

We could find no general relationship between maximal activity of a synthetic promoter and sensitivity. If core promoters do not influence sensitivity, we predicted that changing the core promoters of native (full-length) promoters should have no effect on sensitivity. To directly test if this was the case, we constructed strains with the full-length *astC* and *gadB* promoters, but the core promoter swapped. *astC* and *gadB* were chosen because they are strongly sensitive and insensitive, respectively. These constructs started with previously studied *lacZ* fusions driven by the regions upstream of *astC* and *gadB* (Wong et al., 2017). These fusions contained bases approximately −450 to +170 relative to the transcription start site, including a single annotated core promoter and transcription start site, and all known transcription factor binding sites. We then used mutagenesis to swap the core promoters (i.e. switch the bases of the core promoter of *gadB* with the bases of the *astC* core promoter in the full-length *gadB* promoter, and vice versa.) These core promoter swaps had a negligible effect on sensitivity (Figure 3), although *astC* core into *gadB* had a large effect on total activity. The *astC* core into *gadB* was insensitive, as was the native *gadB*. These two promoters differ slightly, though significantly, in sensitivity (p = 0.023, two-sample randomization test, 100,000 replicates). The *gadB* core swapped into *astC* is sensitive, just like the native *astC*. They do not differ significantly in sensitivity (p = 0.41, two-sample randomization test, 100,000 replicates).

## DISCUSSION

Changes to core promoter sequences did not alter sensitivity to RpoS, an unexpected finding. While maximal gene expression induced by RpoS is clearly affected by the weaker binding sites tested here, the shape of the response to increasing RpoS concentrations remained largely unaffected across the twelve constructs tested here (Figure 1), and there is no correlation between maximum strength and sensitivity. Our promoter swap experiments further show that RpoS-DNA interactions do not determine sensitivity, as the full-length promoters retain their pattern of sensitivity even when the core promoter is replaced with one from a promoter showing a very different pattern. Taken together, our results suggest that sensitivity of a promoter is controlled by factors outside of the core promoter.

Our findings that sensitivity is controlled by interactions other than those between a DNA binding protein and the DNA its binds place our work in contrast to other studied examples, including the sigma factor Spo0A in *B. subtilis* and the transcription factors PhoB and LexA in *E. coli* and CodY in *B. subtilis* (Brinsmade et al., 2014; Culyba et al., 2018; Fujita et al., 2005; Gao and Stock, 2015). Our findings are consistent with previous bioinformatic work demonstrating that there is no sequence motif that distinguishes sensitive from insensitive promoters (Wong et al., 2017). In addition, the finding that a subset of transcription factors are enriched for binding either sensitive or insensitive promoters is consistent with the notion that interactions outside the core promoter determine sensitivity (Wong et al., 2017). Finally, insensitive patterns of transcription cannot be explained by simple Michaelis-Menten kinetics of interacting core promoters and RpoS, also consistent with other regulation driving the response.

The work reported here suggests that because the sensitivity of a promoter and its maximal strength are not coupled, then they can be altered independently, either by evolution or by synthetic biologists. As genes with sensitive and insensitive responses differ in their biological functions, it seems that these expression profiles serve important roles in the timing of gene expression and responses to different stresses (Wong et al., 2017). Our results suggest that this behavior is not mediated by variation in the core promoter, and instead implicates the need for additional regulation by transcription factors to achieve the coordinated timing of transcriptional responses to changing RpoS levels.

## ACKNOWLEDGEMENTS

Thanks to Roya Amini-Naieni, Lakshmi Batachari, Carla Becker, Moira Dillon, and Asaul Gonzalez for technical help, and to Jane Liu and Pete Chandrangsu for comments on an earlier version of the manuscript. This work was supported by NSF Grant #1716794 to Dan Stoebel and HHMI Undergraduate Science Education award #52007544 to Harvey Mudd College.

## Supplemental material

**Table S1.**
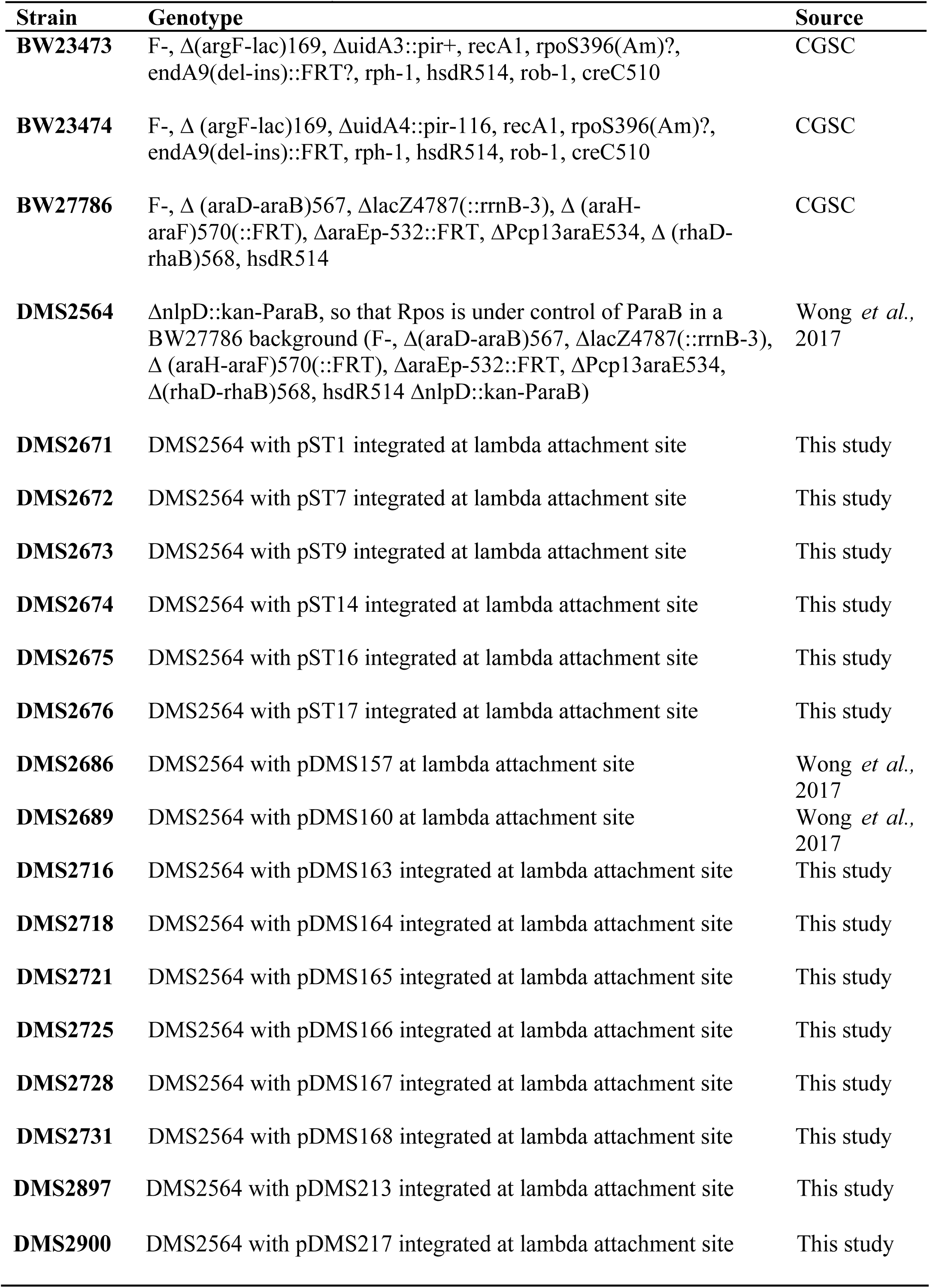
Strains used in this study.

| Strain | Genotype | Source |
| --- | --- | --- |
| **BW23473** | F-, $\Delta(\text{argF-lac})169$, $\Delta\text{uidA3::pir+}$, $\text{recA1}$, $\text{rpoS396(Am)?}$, $\text{endA9(del-ins)::FRT?}$, $\text{rph-1}$, $\text{hsdR514}$, $\text{rob-1}$, $\text{creC510}$ | CGSC |
| **BW23474** | F-, $\Delta(\text{argF-lac})169$, $\Delta\text{uidA4::pir-116}$, $\text{recA1}$, $\text{rpoS396(Am)?}$, $\text{endA9(del-ins)::FRT}$, $\text{rph-1}$, $\text{hsdR514}$, $\text{rob-1}$, $\text{creC510}$ | CGSC |
| **BW27786** | F-, $\Delta(\text{araD-araB})567$, $\Delta\text{lacZ4787(::rrnB-3)}$, $\Delta(\text{araH-araF})570(::\text{FRT})$, $\Delta\text{araEp-532::FRT}$, $\Delta\text{Pcp13araE534}$, $\Delta(\text{rhaD-rhaB})568$, $\text{hsdR514}$ | CGSC |
| **DMS2564** | $\Delta\text{nlpD::kan-ParaB}$, so that Rpos is under control of ParaB in a BW27786 background (F-, $\Delta(\text{araD-araB})567$, $\Delta\text{lacZ4787(::rrnB-3)}$, $\Delta(\text{araH-araF})570(::\text{FRT})$, $\Delta\text{araEp-532::FRT}$, $\Delta\text{Pcp13araE534}$, $\Delta(\text{rhaD-rhaB})568$, $\text{hsdR514}$ $\Delta\text{nlpD::kan-ParaB}$) | Wong *et al.*, 2017 |
| **DMS2671** | DMS2564 with pST1 integrated at lambda attachment site | This study |
| **DMS2672** | DMS2564 with pST7 integrated at lambda attachment site | This study |
| **DMS2673** | DMS2564 with pST9 integrated at lambda attachment site | This study |
| **DMS2674** | DMS2564 with pST14 integrated at lambda attachment site | This study |
| **DMS2675** | DMS2564 with pST16 integrated at lambda attachment site | This study |
| **DMS2676** | DMS2564 with pST17 integrated at lambda attachment site | This study |
| **DMS2686** | DMS2564 with pDMS157 at lambda attachment site | Wong *et al.*, 2017 |
| **DMS2689** | DMS2564 with pDMS160 at lambda attachment site | Wong *et al.*, 2017 |
| **DMS2716** | DMS2564 with pDMS163 integrated at lambda attachment site | This study |
| **DMS2718** | DMS2564 with pDMS164 integrated at lambda attachment site | This study |
| **DMS2721** | DMS2564 with pDMS165 integrated at lambda attachment site | This study |
| **DMS2725** | DMS2564 with pDMS166 integrated at lambda attachment site | This study |
| **DMS2728** | DMS2564 with pDMS167 integrated at lambda attachment site | This study |
| **DMS2731** | DMS2564 with pDMS168 integrated at lambda attachment site | This study |
| **DMS2897** | DMS2564 with pDMS213 integrated at lambda attachment site | This study |
| **DMS2900** | DMS2564 with pDMS217 integrated at lambda attachment site | This study |

**Table 2.**
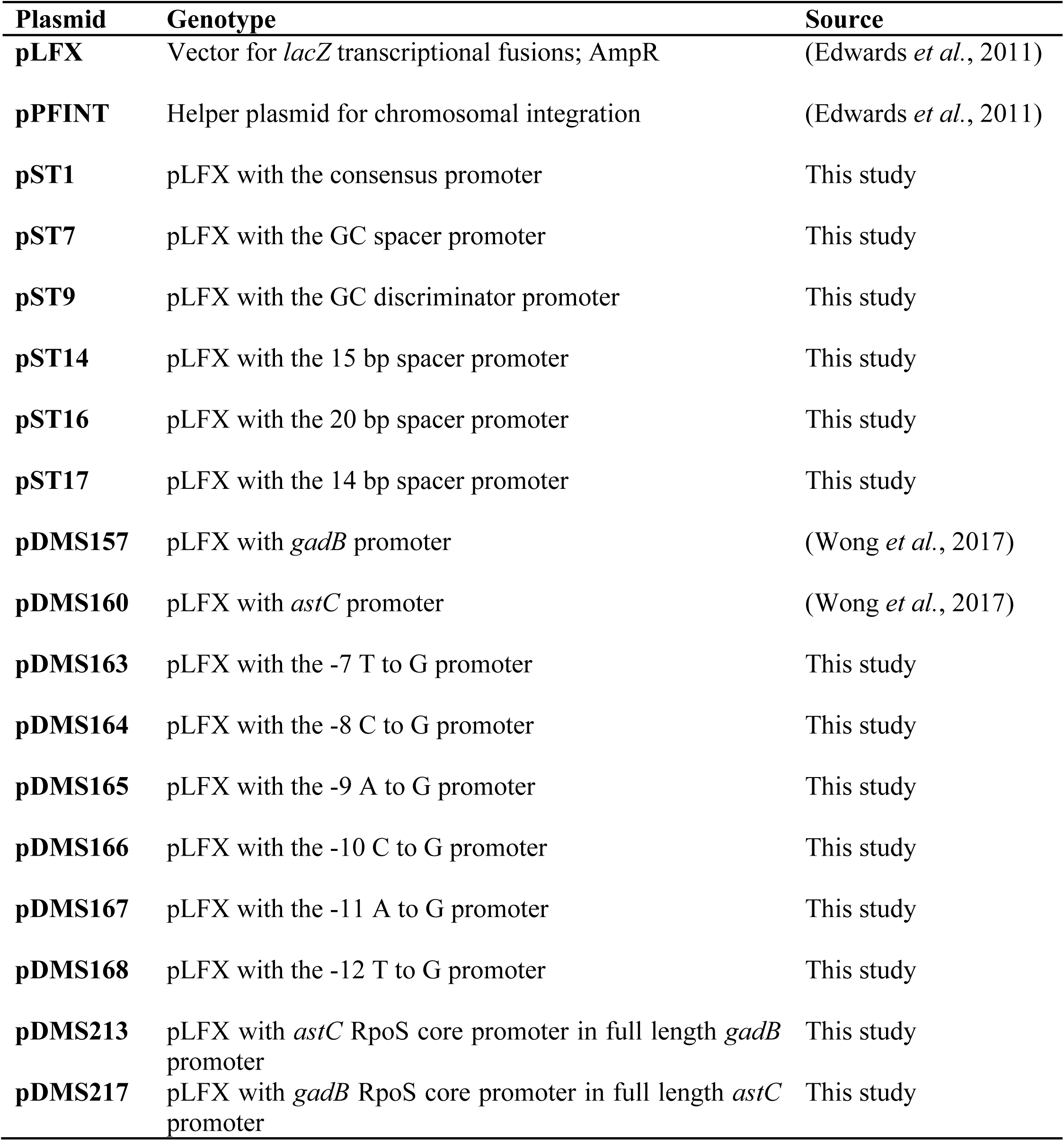
Plasmids used in this study.

| **Plasmid** | **Genotype** | **Source** |
| --- | --- | --- |
| **pLFX** | Vector for *lacZ* transcriptional fusions; AmpR | (Edwards *et al.*, 2011) |
| **pPFINT** | Helper plasmid for chromosomal integration | (Edwards *et al.*, 2011) |
| **pST1** | pLFX with the consensus promoter | This study |
| **pST7** | pLFX with the GC spacer promoter | This study |
| **pST9** | pLFX with the GC discriminator promoter | This study |
| **pST14** | pLFX with the 15 bp spacer promoter | This study |
| **pST16** | pLFX with the 20 bp spacer promoter | This study |
| **pST17** | pLFX with the 14 bp spacer promoter | This study |
| **pDMS157** | pLFX with *gadB* promoter | (Wong *et al.*, 2017) |
| **pDMS160** | pLFX with *astC* promoter | (Wong *et al.*, 2017) |
| **pDMS163** | pLFX with the -7 T to G promoter | This study |
| **pDMS164** | pLFX with the -8 C to G promoter | This study |
| **pDMS165** | pLFX with the -9 A to G promoter | This study |
| **pDMS166** | pLFX with the -10 C to G promoter | This study |
| **pDMS167** | pLFX with the -11 A to G promoter | This study |
| **pDMS168** | pLFX with the -12 T to G promoter | This study |
| **pDMS213** | pLFX with *astC* RpoS core promoter in full length *gadB* promoter | This study |
| **pDMS217** | pLFX with *gadB* RpoS core promoter in full length *astC* promoter | This study |

**Table 3.**
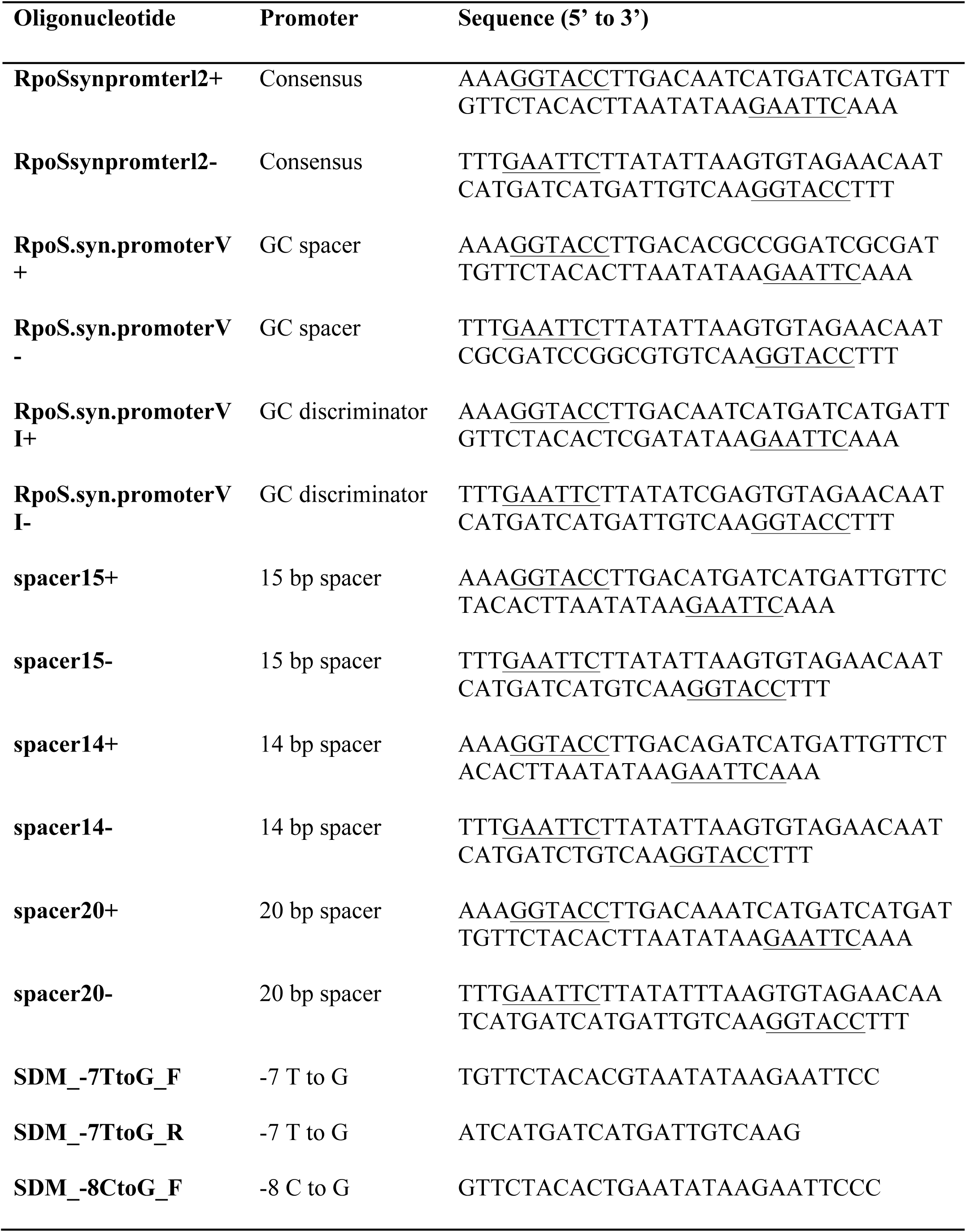
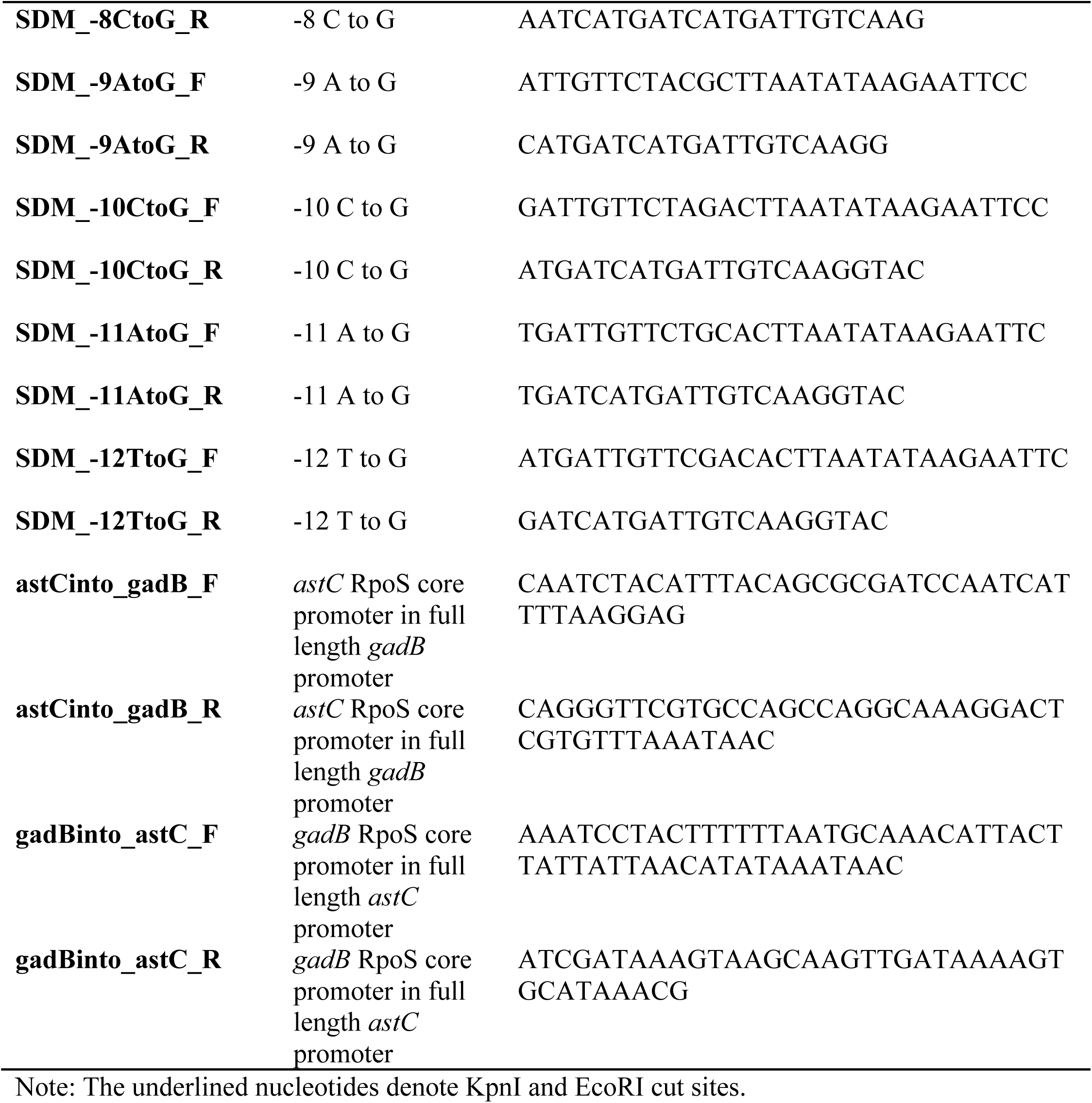
Oligonucleotides used for creating promoters.

